# Azithromycin reduces systemic inflammation and provides survival benefit in murine model of polymicrobial sepsis

**DOI:** 10.1101/303610

**Authors:** Anasuya Patel, Jiji Joseph, Hariharan Periasamy, Santosh Mokale

## Abstract

Sepsis is a life threatening systemic inflammatory condition triggered as a result of excessive host immune response to infection. In the past, drugs modulating immune reactions have demonstrated protective effect in sepsis. Azithromycin (macrolide antibiotic) with immunomodulatory activity was therefore evaluated in combination with ceftriaxone in a more clinically relevant murine model of sepsis induced by caecal ligation and puncture (CLP). First, mice underwent CLP and 3 h later were administered with vehicle, sub-effective dose of ceftriaxone (100 mg/kg, *subcutaneous*) alone or in combination with immunomodulatory dose of azithromycin (100 mg/kg, *intraperitoneal*). Survival was then monitored for 5 days. Parameters like body temperature, blood glucose, total white blood cell count, plasma glutathione (GSH), plasma and lung myeloperoxidase (MPO) as well as cytokine (interleukin IL-6, IL-1β, tumor necrosis factor-α) levels along with bacterial load in blood, peritoneal fluid and lung homogenate were measured 18 h after CLP challenge. Combination group significantly improved the survival of CLP mice. It attenuated the elevated levels of inflammatory cytokines and MPO in plasma and lung tissue and increased the body temperature, blood glucose and GSH which were otherwise markedly decreased in CLP mice. Ceftriaxone exhibited significant reduction of bacterial count in blood, peritoneal fluid and lung homogenate, while co-administration of azithromycin did not further reduce it. This confirms that survival benefit by azithromycin was due to immunomodulation and not by its antibacterial action. Findings of this study indicate that azithromycin in combination with ceftriaxone could exhibit clinical benefit in sepsis.

## INTRODUCTION

Sepsis is a generalized systemic inflammatory condition elicited by pro-inflammatory cytokines released by host immune cell in response to exotoxin or endotoxins secreted by bacteria. Overproduction of these cytokines is associated with multiple organ failure which is the major cause of death in critically ill and elderly patients. Despite recent advances in our understanding of the pathophysiological mechanism of sepsis and improved antimicrobial therapy, the mortality rate from sepsis remains frustratingly high. Unfortunately many of the therapeutic options proposed over the years for the management of sepsis and its complication have either failed to meet their initial expectations or remained unproved. Strategies are currently being developed to minimize the inflammatory response associated with sepsis through immunotherapy (1).

Several reports have provided evidence of immunomodulatory activity of macrolide antibiotics (2, 3, 4). As a result, macrolides have proved beneficial in chronic pulmonary inflammatory conditions like diffuse panbronchiolitis, cystic fibrosis, asthma and bronchiectasis (5, 6). Due to this behaviour, they have earlier demonstrated clinical benefits in critically ill patients with CABP or Gram negative sepsis by reducing the disease severity, length of hospital stay and mortality rates (7, 8).

Azithromycin, belonging to macrolide antibacterial class has been earlier reported to provide survival benefit in the lipopolysaccharide (LPS) induced sepsis model (9). In the present study, we aimed to extend this investigation in the caecal ligation and puncture (CLP) model of polymicrobial sepsis. CLP model is widely preferred sepsis model as it closely mimics the clinical condition of human sepsis (10). Azithromycin acts on Gram positive bacteria and is bacteriostatic, therefore in this study it was combined with ceftriaxone, a third generation cephalosporin antibiotic to control the progression of polymicrobial infection.

## RESULTS

### Survival in Sepsis model

In the LPS induced sepsis model, azithromycin administered 1 h prior to lethal dose of LPS showed dose dependent improvement in 24 h survival. Azithromycin at 100 mg/kg, *i.p*. provided survival in 75 % of mice as compared to 12.5% in LPS group, which was statistically significant (p<0.05) **(Figure 1)**. Survival benefit of azithromycin was further determined in CLP sepsis, a model having best correlation to human sepsis. In this model, azithromycin at 100 mg/kg dose provided protection in 37.5 % of mice in contrast to 75% in LPS treated mice, indicating that azithromycin might have little or no effect on the spread of infection. In order to control bacterial infection, azithromycin was further combined with ceftriaxone (broad spectrum antibiotic) at sub-effective dose. This dose of ceftriaxone was identified from our initial survival studies in CLP mice where it provided protection in 37.5 % of mice (data not shown). Subsequently, sub-effective dose of ceftriaxone (100 mg/kg, *s.c*.) was combined with immunomodulatory dose of azithromycin (100 mg/kg, *i.p*.) and the survival was monitored for 5 days. A log-rank test analysis of the 5 day survival curve in CLP sepsis showed that survival rate in combination group was significantly higher compared to CLP control (p<0.01) **(Figure 2)**.

**Figure 1.**
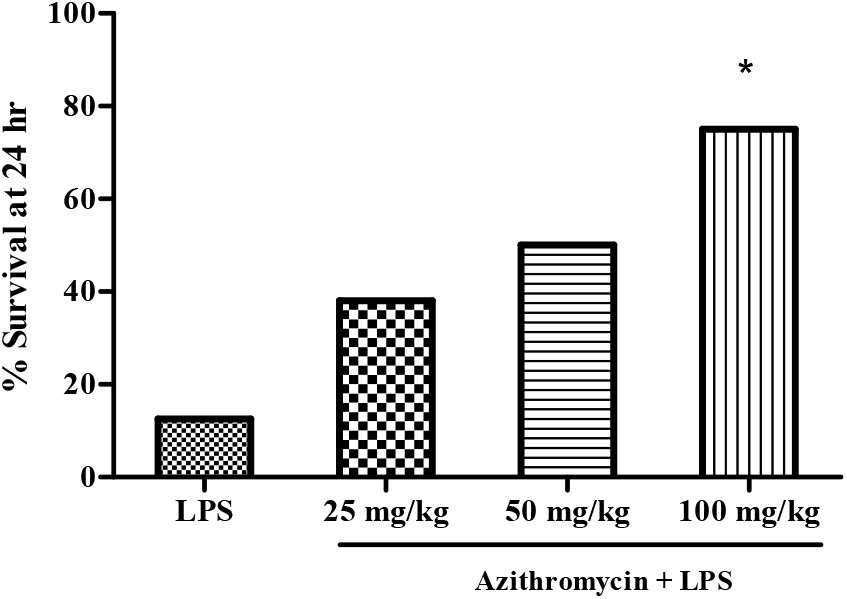
Survival in LPS induced mouse sepsis model. Vehicle (saline) and azithromycin (25, 50, 100 mg/kg) was administered intraperitoneally to mice (n=8), 1 h prior to LPS treatment (1500 µg/mouse) and 24 h survival was monitored. In LPS group only 12.5 % of mice survived whereas LPS treated mice administered azithromycin exhibited dose dependent improvement in survival with 100 mg/kg dose providing protection in 75% of mice.

**Figure 2.**
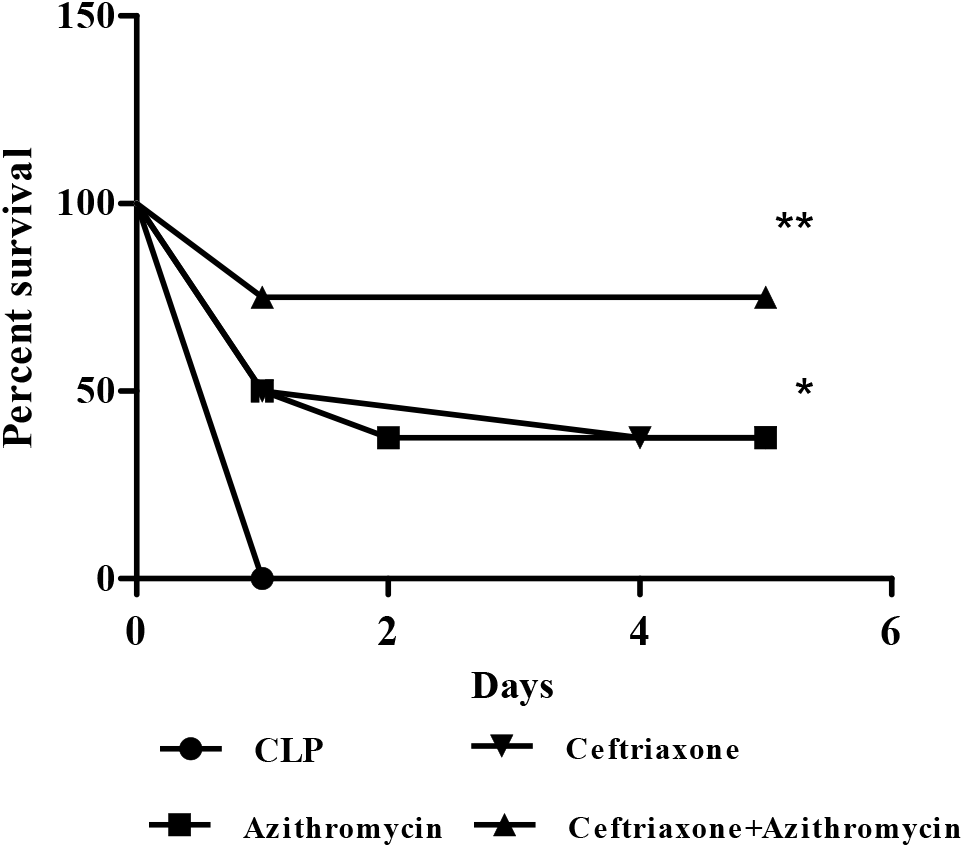
Survival data in CLP mice. CLP challenged mice (n=8) were administered ceftriaxone (100 mg/kg, *s.c*.), azithromycin (100 mg/kg, *i.p*.), both in combination and survival was monitored for 5 days. Ceftriaxone and azithromycin provided protection in 37.5 % of mice while combination provided survival in 75% of mice.

### Effect on blood glucose, body temperature and total WBC count

Blood glucose, body temperature and WBC count are affected during systemic inflammation. WBC count was significantly (p<0.001) reduced in LPS group (2.77×10^3^/µL ± 0.14) compare to normal group (7.02×10^3^/µL ± 0.63) which was not reversed by treatment with azithromycin (**Figure 3A**). Blood glucose and body temperature were also significantly reduced in LPS treated mice. The values of blood glucose and body temperature in normal mice were 119.5 ± 2.4 mg/dL and 99.2 ± 0.7 ºF which dropped to 20.67 ± 2.1 mg/dL (p>0.001) and 92.0 ± 0.7 ºF (p<0.001), respectively in LPS treated mice. Azithromycin at 100 mg/kg, *i.p*., significantly elevated the blood glucose (51.5 ± 9.0 mg/dL, p<0.01) and body temperature (96.28 ± 0.4 ºF, p<0.001) (**Figure 3B and 3C**).

**Figure 3.**
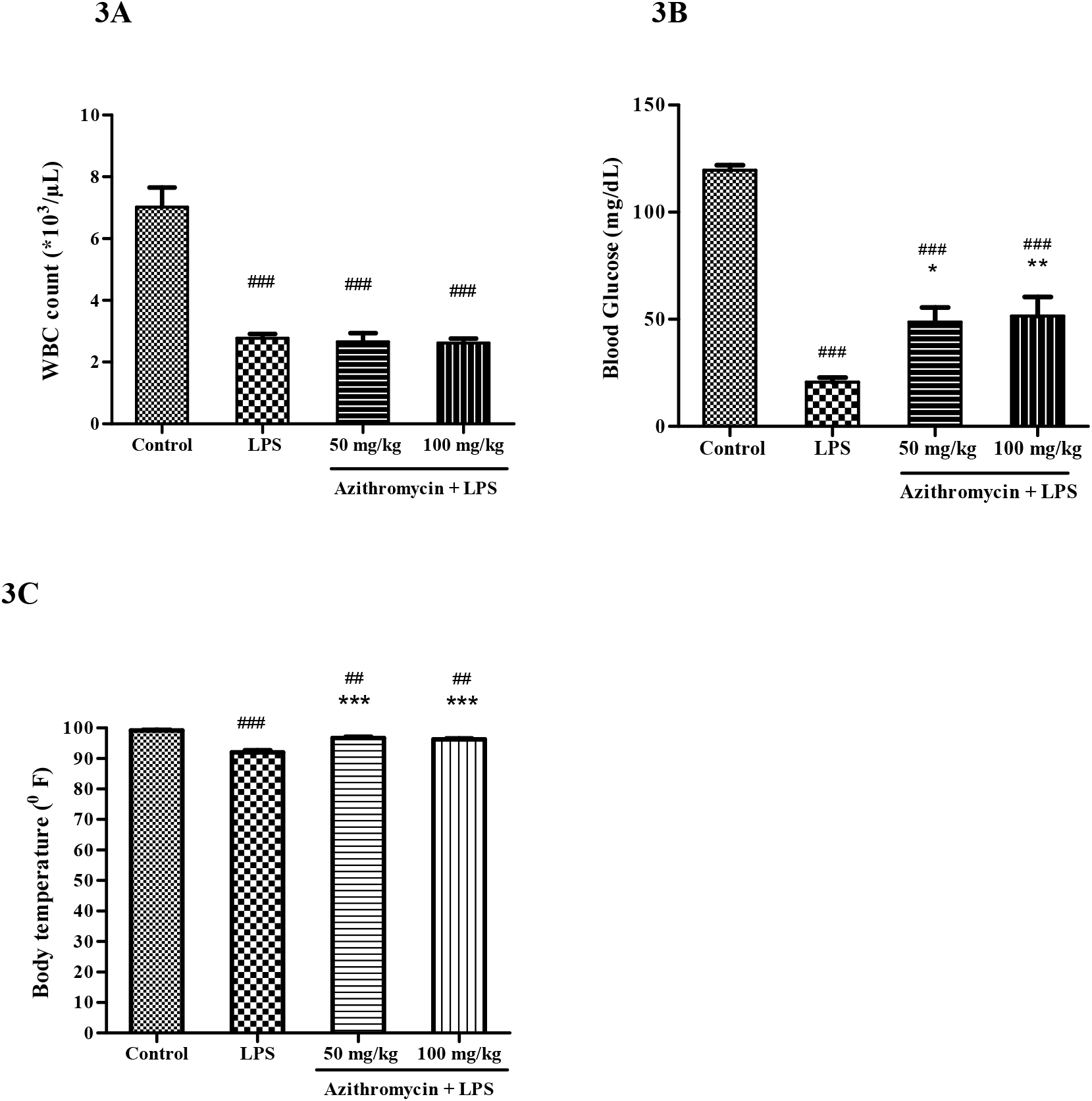
Effect on WBC (A), blood glucose (B) and body temperature (C) in LPS treated mice: Mice (n=8) were administered azithromycin (50 and 100 mg/kg, *i.p*.) or vehicle (saline), 1 h prior to LPS treatment (1000 µg/mouse) and the body temperature, blood glucose and WBC count were determined at 18 h after LPS injection in 6 mice. LPS group showed significantly lower WBC, blood glucose and body temperature compared to normal control group. In LPS treated mice, azithromycin exhibited no effect on the WBC count; while blood glucose and body temperature increased significantly at both the doses compare to LPS group. Values represent means ±SEM. ##p> 0.01, ###p>0.001 indicated value versus control; *p> 0.05, **p>0.01, ***p>0.001 in LPS plus azithromycin versus LPS group

In the CLP mice also, the total WBC count was significantly lower than the sham control group (1.13×10^3^/µL ± 0.14 versus 6.18 ×10^3^/µL ± 0.25; p<0.001). Ceftriaxone treatment increased WBC count although not statistically significant (2.77×10^3^/µL ± 0.73). While, azithromycin combined with ceftriaxone neither increased nor decreased the WBC count (2.58×10^3^/µL ± 0.36) (**Figure 4A**). In comparison to the sham group, mice with CLP caused significant reduction in the blood glucose (from 99.55 ± 3.3 to 29.0 ± 13.0 mg/dL; p<0.001) and body temperature (from 99.33 ± 0.4 to 92.97 ± 1.1ºF; p<0.001). CLP mice treated with ceftriaxone (100 mg/kg, *s.c*.) significantly increased the blood glucose (54.0 ± 7.6 mg/dL; p<0.05) as well as body temperature (96.33 ± 0.4ºF; p<0.01), and in combination with azithromycin there was further increase in the blood glucose (72.7 ± 6.5 mg/dL; p<0.01) and body temperature (97.0 ± 0.5 ºF; p<0.01) (**Figure 4B and 4C**).

**Figure 4.**
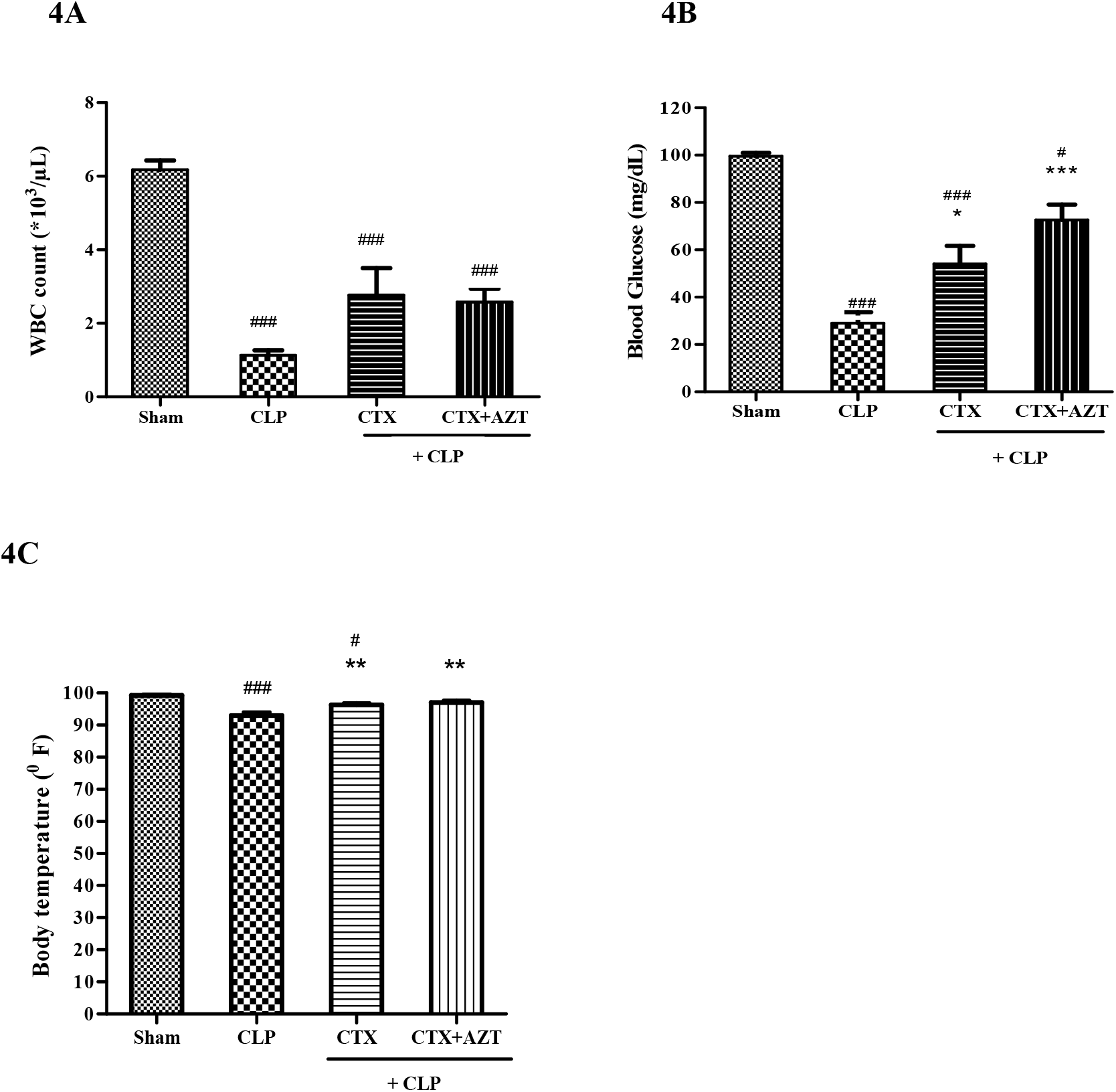
Effect of ceftriaxone (CTX) and ceftriaxone (CTX) + azithromycin (AZT) on WBC (A), blood glucose (B) and body temperature (C) in CLP mice: After 3 h of CLP challenge, mice (n=8) were administered vehicle (saline), ceftriaxone (100 mg/kg, *s.c*.) and ceftriaxone (100 mg/kg, *s.c*.) + azithromycin (100 mg/kg, *i.p*.) and 18 h later WBC, blood glucose and body temperature were measured. CLP mice significantly reduced WBC, blood glucose and body temperature versus sham group. An increase in WBC was observed in ceftriaxone and combination group; however it was not statistically significant versus CLP control. There was a significant increase in blood glucose and body temperature in ceftriaxone and combination group; however the improvement in blood glucose and body temperature was better in combination. Values represent means ±SEM. #p<0.05, ###p<0.001 indicated value versus control; *p>0.05, **p>0.01 and ***p> 0.001 indicates CLP plus ceftriaxone or CLP plus (ceftriaxone + azithromycin) versus CLP group.

### Effect on GSH and MPO levels

The levels of GSH (an endogenous thiol antioxidant) and MPO (an inflammatory enzyme) are considerably altered during sepsis. There was a significant suppression in plasma GSH level and significant elevation in plasma MPO levels in LPS treated mice compared to normal group. GSH decreased from 2.37 ± 0.2 to 0.82 ± 0.04 mg/mL (p<0.001) and MPO increased from 290.2 ± 88.9 to 871.81 ± 86.2 µM/min/mL (p< 0.001), while LPS treated mice administered azithromycin produced dose dependent rise in GSH (0.96 ± 0.1 mg/mL at

50 mg/kg and 1.04 ± 0.03 mg/mL at 100 mg/kg) and decrease in MPO levels (694.03 ± 67.7 µM/min/mL at 50 mg/kg and 629.08 ± 42.3 µM/min/mL at 100 mg/kg), however the effect was significant (p<0.05) only at 100 mg/kg **(Figure 5A and 5B)**.

**Figure 5.**
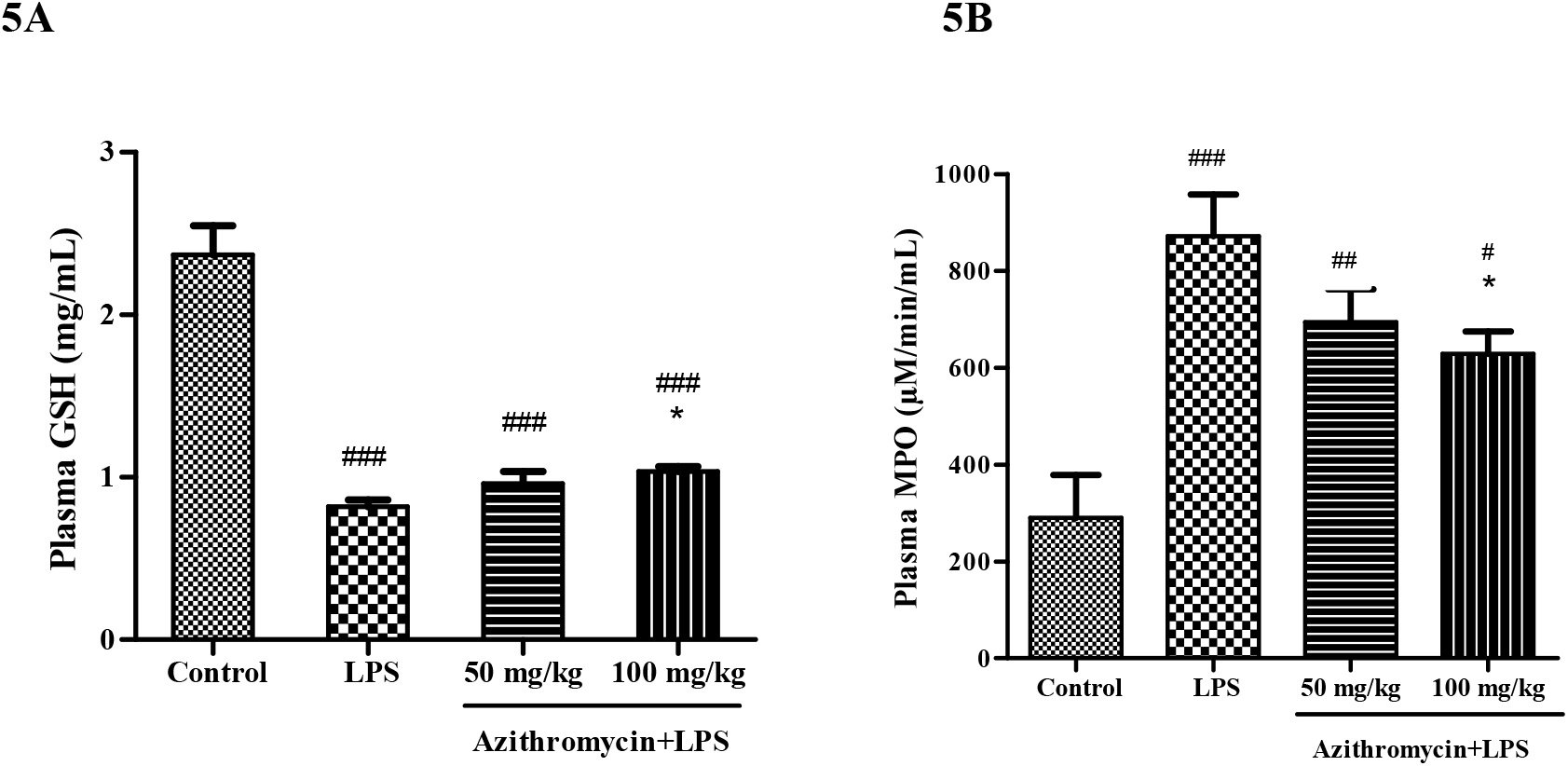
Effect on GSH (A) and MPO (B) in the LPS treated mice: Mice (n=8) were administered vehicle (saline) and azithromycin (50 and 100 mg/kg, *i.p*.), 1 h prior to LPS challenge and the plasma GSH and MPO were measured at 18 h post LPS injection in 6 mice. LPS group showed significant decrease in GSH and increase in MPO levels compared to control group. While, LPS treated mice administered azithromycin (100 mg/kg), demonstrated significant increase in GSH and decrease in MPO. Values represent means ±SEM. #p>0.05, ##p> 0.01, ###p>0.001 indicated value versus control; *p> 0.05 indicates LPS plus azithromycin versus LPS group.

In the CLP challenged mice, plasma GSH was significantly reduced and plasma MPO was significantly increased compared to sham group (GSH: 1.02 ± 0.1 in CLP versus 2.08 ± 0.2 mg/mL in sham group, p<0.01 and MPO: 620.64 ± 45.9 in CLP versus 105.84 ± 33.8 µM/min/mL in sham group, p<0.01). Ceftriaxone treatment did not cause significant change in the GSH and MPO. While, combination group significantly elevated GSH (1.98 ± 0.3 mg/mL; p<0.01) and decreased MPO (275.13 ± 50.4 µM/min/mL; p< 0.01) **(Figure 6A and 6B)**. In lung tissue, MPO levels in CLP mice increased significantly compared to sham group (60.27 Vs 8.51 U/gm of tissue). Ceftriaxone did not have any effect on lung MPO (52.04 U/gm), while ceftriaxone and azithromycin combination significantly reduced its value (44.20 U/gm) (**Figure 6C**).

**Figure 6.**
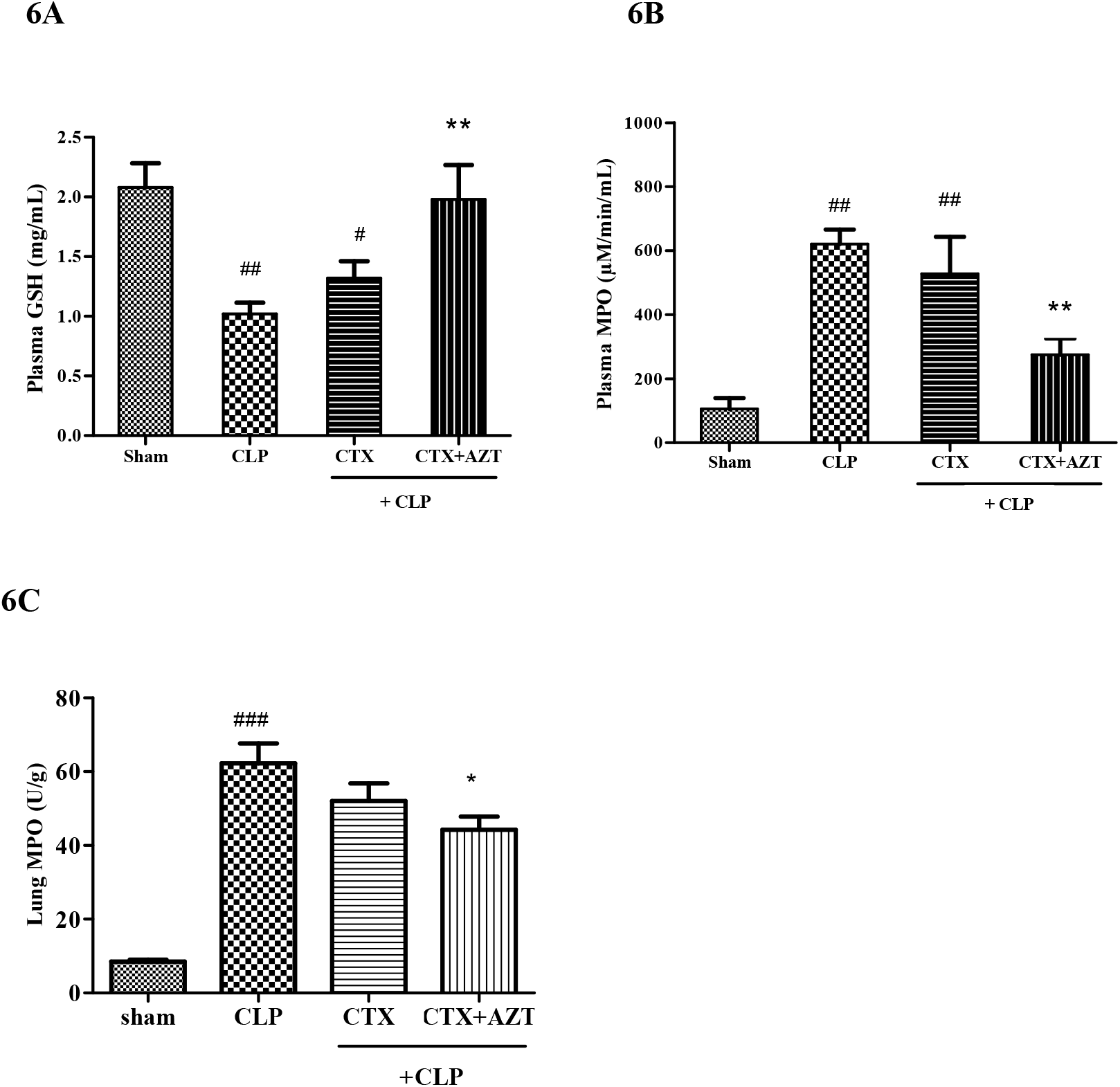
Effect of ceftriaxone (CTX) and ceftriaxone (CTX) + azithromycin (AZT) on GSH (A) and MPO (B) in CLP mice: Vehicle (saline), ceftriaxone (100 mg/kg, *s.c*.) and ceftriaxone (100 mg/kg, *s.c*.) + azithromycin (100 mg/kg, *i.p*.) were administered 3 h after CLP (n=8). Plasma GSH, plasma and lung MPO were measured after 18 h of CLP. CLP mice significantly reduced GSH and increased MPO versus sham group. CLP mice treated with ceftriaxone had no effect on GSH and MPO but the combination group significantly elevated plasma GSH and diminished plasma and lung MPO levels versus CLP group. Values represent means ±SEM. #p<0.05, ## p<0.01 indicated value versus sham group; *p> 0.05 and **p>0.01 indicates CLP plus (ceftriaxone + azithromycin) versus CLP group

### Effect on circulating cytokine levels

Plasma cytokine concentrations in LPS sepsis model are illustrated in **figure 7**. Proinflammatory cytokines are the main mediators of inflammation during sepsis. As expected, significant increase in IL-6, IL-1β and TNF-α levels occurred in the LPS treated group compared to control group. The values of IL-6, IL-1β and TNF-α in normal control group were 18.47 ± 3.5, 8.28 ± 0.6 and 11.15 ± 0.5 pg/mL which significantly increased to 344.44 ± 20.8 (p>0.001), 983.33 ± 72.7 (p<0.001) and 113 ± 12.4 pg/mL (p<0.001), respectively in LPS treated mice. Azithromycin administered to LPS challenged mice showed dose dependent inhibition of these cytokines. At 100 mg/kg, azithromycin showed distinct reduction in IL-6 (47.28 ± 8.4 pg/mL; p<0.001), IL-1β (15 ± 1.2 pg/mL; p<0.001) and TNF-α (34.63 ± 1.3 pg/mL; p<0.001) compared to LPS group.

**Figure 7.**
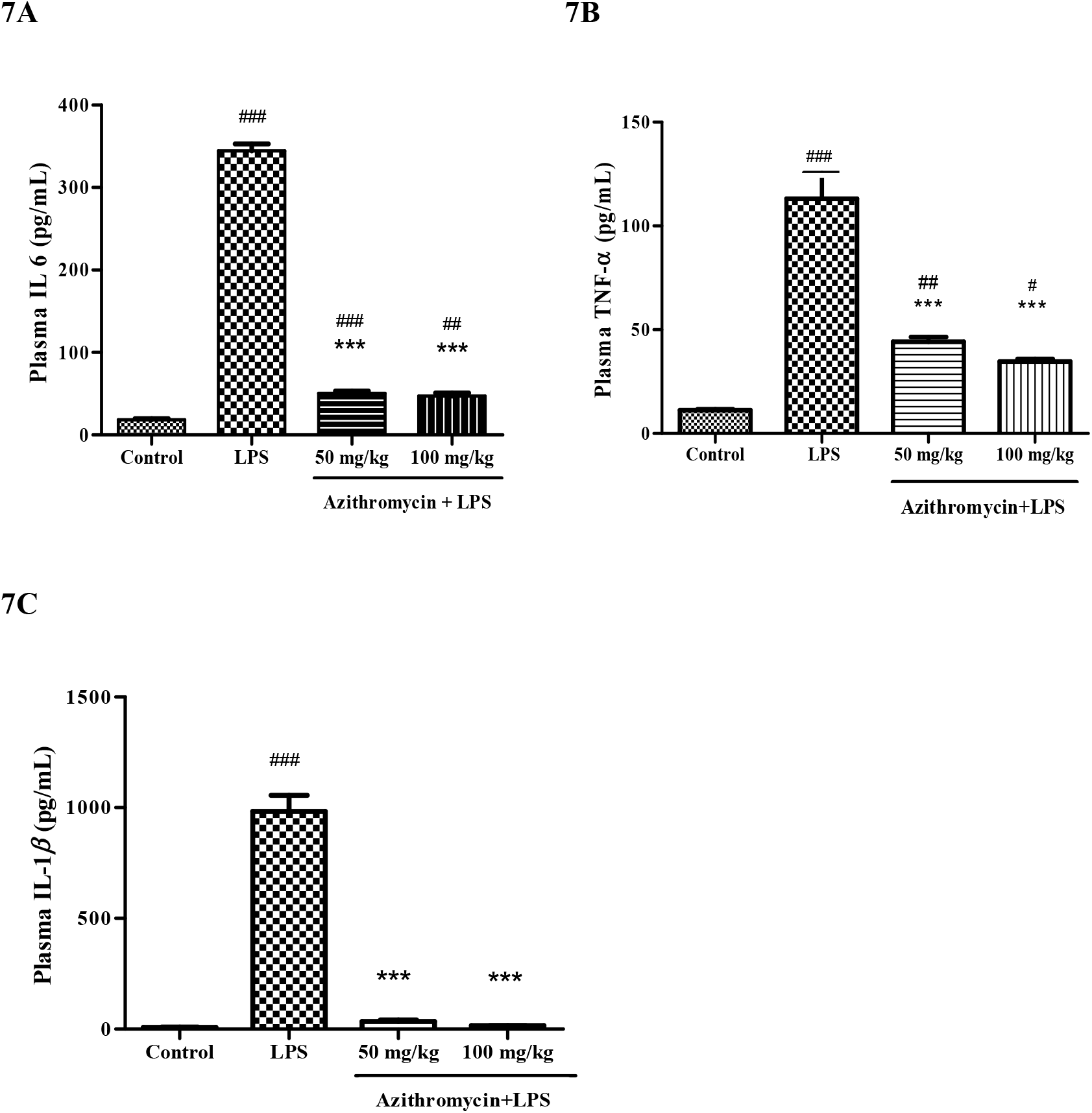
Effect on plasma cytokines IL-6 (A), TNF-α (B) and IL-1β (C) in the LPS treated mice: Mice (n=8) were administered vehicle (saline) and azithromycin (50 and 100 mg/kg, *i.p*.), 1 h prior to LPS challenge and the plasma cytokines were measured at 18 h after LPS injection in 6 mice. LPS group showed significant increase in these cytokines compared to normal control. While, LPS treated mice administered azithromycin (50 and 100 mg/kg), demonstrated significant decrease in IL-6, IL-1β and TNF-α. Values represent means ±SEM. #p>0.05, ##p> 0.01, ###p>0.001 indicated value versus normal control; ***p> 0.001 indicates LPS plus azithromycin versus LPS group.

Similarly, CLP challenged mice showed statistically significant rise in the plasma cytokine levels compared to sham group (IL-6, TNF-α and IL-1β: 15892 ± 1250.6, 119.3 ± 17.4, 156.61 ± 43.9 pg/mL in CLP group versus 38.92 ± 9.0, 7.7 ± 0.2, 15.76 ± 2.5 pg/mL, respectively in sham group). Ceftriaxone significantly reduced the IL-6 in the CLP mice from 15892 ± 1250.6 to 925 ± 154.5 pg/mL (p<0.001), TNF-α from 119.3 ± 17.4 to 41.03 ±

3.9 pg/mL (p<0.001) and IL-1β from 156.61 ±43.9 to 52.19 ± 6.2 pg/mL (p>0.05). Combination group also demonstrated significant decrease in IL-6, TNF-α and IL-1β compared to CLP group and the values in pg/mL for each were 565 ± 25.8 (p<0.001), 23.10 ± 1.2 (p<0.001) and 36.23 ± 3.4 (p<0.01), respectively (**Figure 8**).

**Figure 8.**
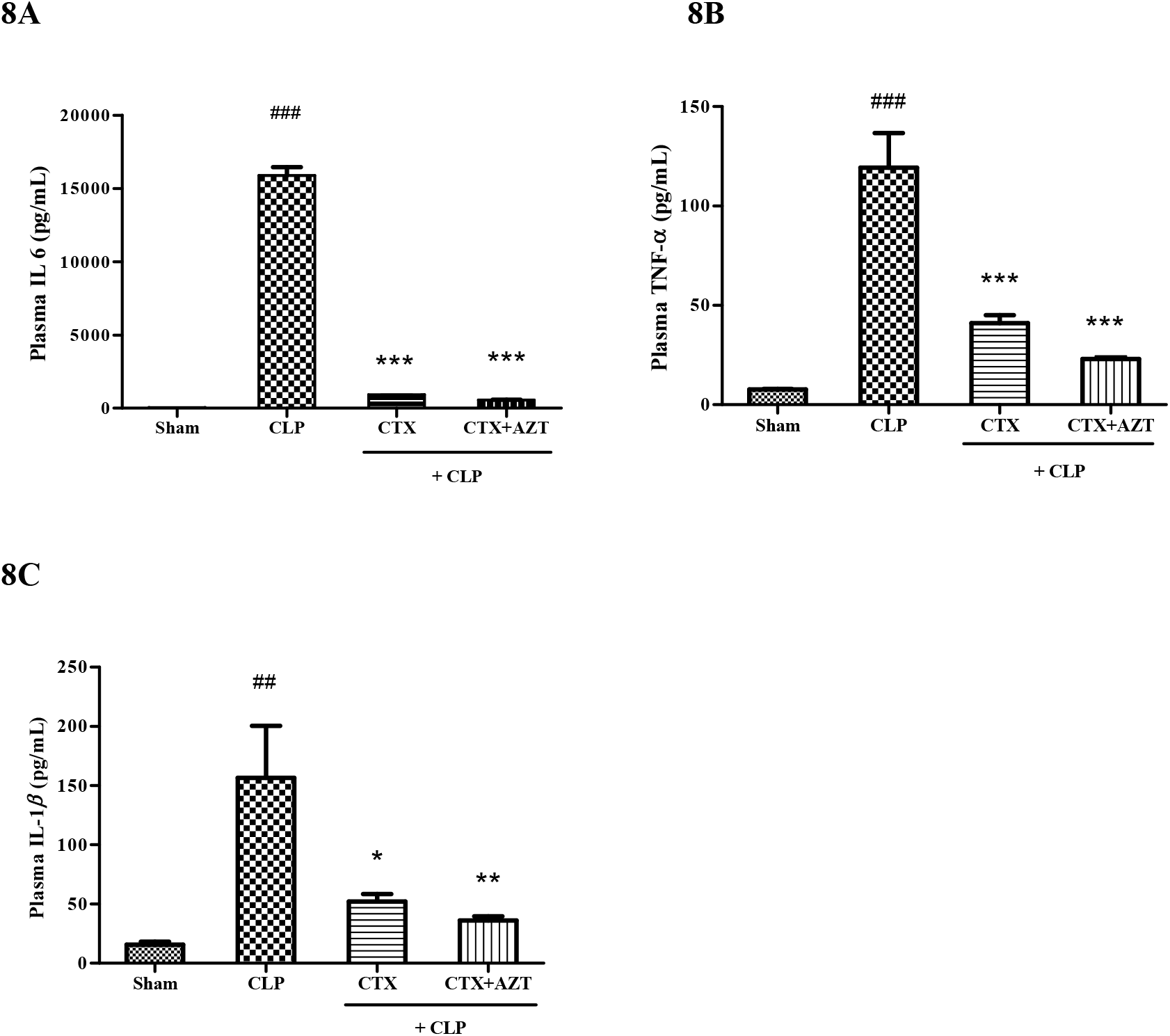
Effect of ceftriaxone (CTX) and ceftriaxone (CTX) + azithromycin (AZT) on Plasma IL-6 (A), TNF-α (B) and IL-1β (C) in the CLP mice: After 3 h of CLP challenge, mice were treated with vehicle, ceftriaxone (100 mg/kg, *s.c*.) and ceftriaxone (100 mg/kg, *s.c*.) + azithromycin (100 mg/kg, *i.p*.) and the plasma cytokine were measured 18 h after CLP (n=6). There was a significant increase in cytokine levels in the CLP mice versus sham group. Ceftriaxone showed significant reduction in elevated cytokine levels, which was further reduced in the combination group. Values represent means ±SEM. ##p>0.01, ###p>0.001 indicated value versus sham group; *p>0.05, **p>0.01 and ***p> 0.001 indicates CLP plus ceftriaxone or CLP plus (ceftriaxone + azithromycin) versus CLP group

Lung levels of IL-6, TNF-α and IL-1β in CLP mice were 466, 251.2 and 526.6 pg/mL, respectively compared to no levels in sham group (data not shown in graph). Ceftriaxone treated CLP mice did not mitigate these cytokine levels while ceftriaxone and azithromycin combination brought about significantly reduction (IL-6: 57.06±4.07, TNF-α: 67.17±13.08 and IL-1β:316.53±21.94 pg/mL) **(Figure 9)**.

**Figure 9.**
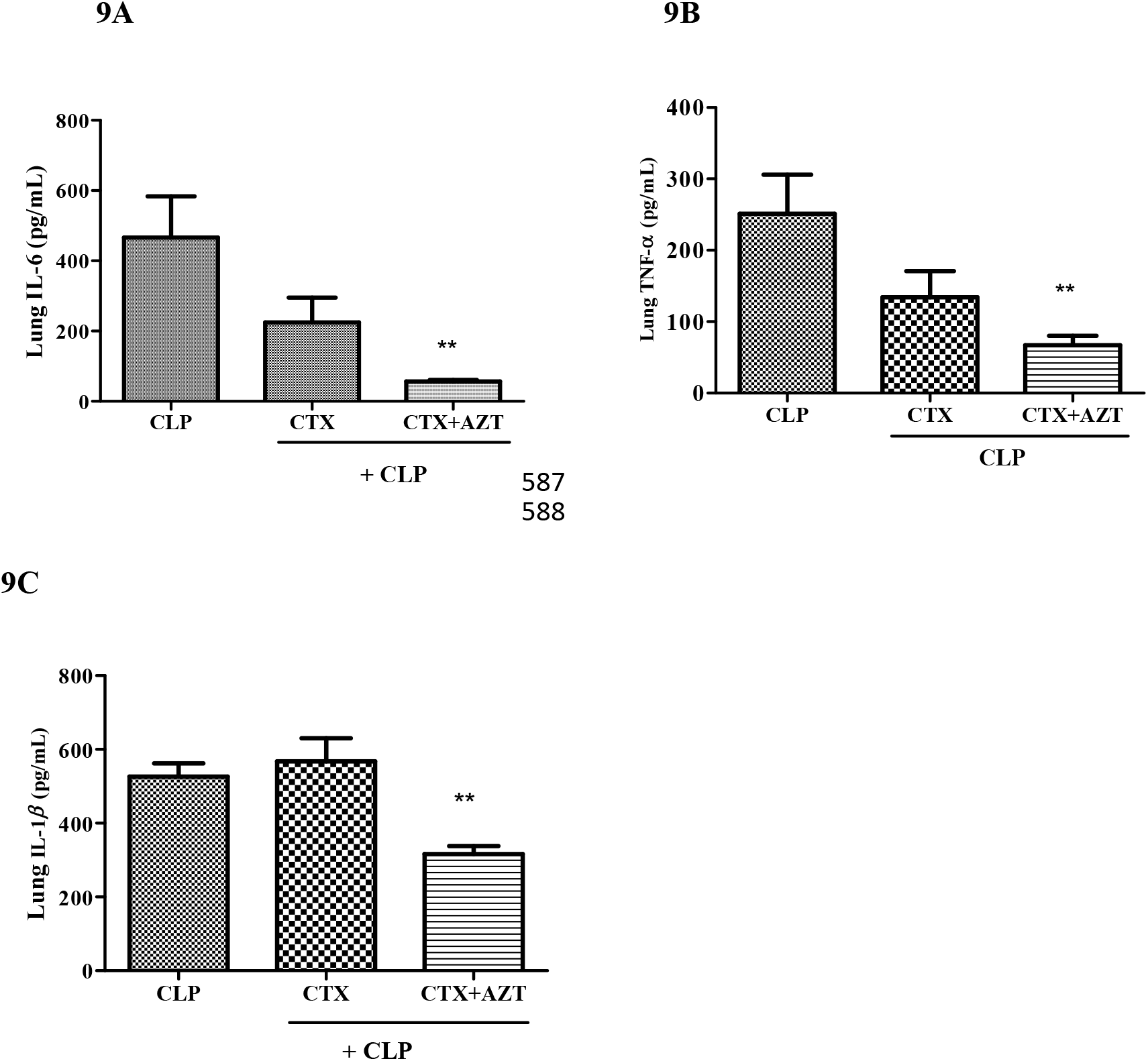
Effect of ceftriaxone (CTX) and ceftriaxone (CTX) + azithromycin (AZT) on Lung IL-6 (A), TNF-α (B) and IL-1β (C) in the CLP mice: After 3 h of CLP challenge, mice were treated with vehicle, ceftriaxone (100 mg/kg, *s.c*.) and ceftriaxone (100 mg/kg, *s.c*.) + azithromycin (100 mg/kg, *i.p*.) and the plasma cytokine were measured 18 h after CLP (n=6). There was a significant increase in cytokine levels in the CLP mice versus undetectable levels in sham group (data not shown). Ceftriaxone did not reduce the elevated cytokine levels in CLP mice while combination group significantly suppressed it. Values represent means ±SEM. **p>0.01 indicates CLP plus (ceftriaxone + azithromycin) versus CLP group

### Effect on Bacterial load in blood, peritoneal fluid and lung homogenate

In order to determine whether the protective effect by combination group in the CLP model was due to synergistic antimicrobial activity, bacterial colony forming units (CFU) was estimated in the blood, peritoneal fluid and lung homogenate following CLP. Ceftriaxone showed significant reduction in bacterial count in blood, lung and peritoneal fluid while azithromycin in combination with ceftriaxone provided no further reduction compared to CLP group. These suggest that azithromycin had no antibacterial effect in this polymicrobial sepsis model **(Figure 10)**.

**Figure 10:**
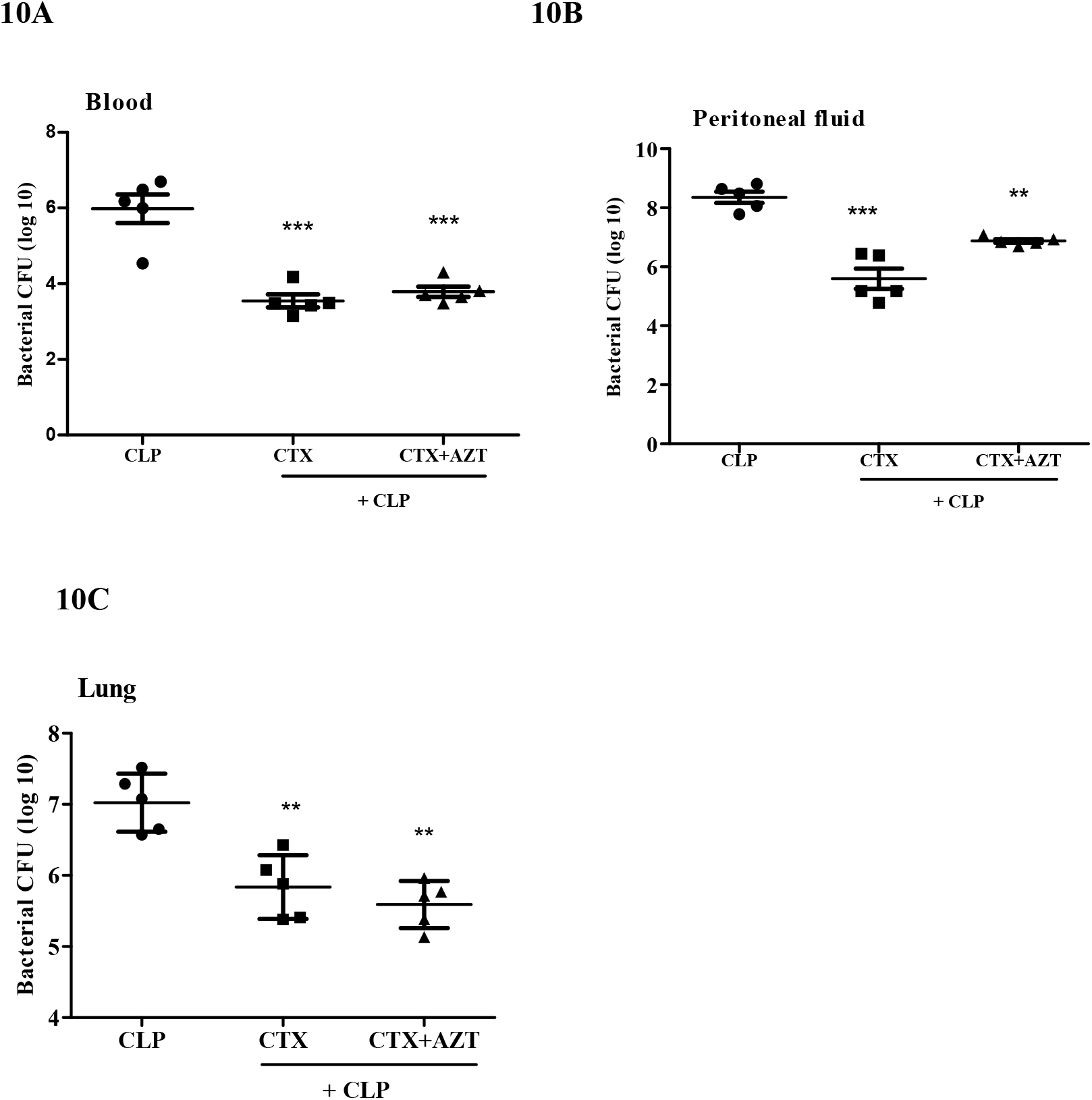
Effect of ceftriaxone (CTX) and ceftriaxone (CTX) + azithromycin (AZT) on bacterial count in blood (A), peritoneal fluid (B) and Lung tissue (C) in CLP mice: Mice (n=6) were treated with ceftriaxone (100 mg/kg, *s.c*.) and ceftriaxone (100 mg/kg, *s.c*.) + azithromycin (100 mg/kg, *i.p*.), 3 h after CLP and the bacterial counting was done at 18 h post CLP challenge in 5 mice. CLP group showed positive bacterial culture while ceftriaxone treatment significantly reduced CFU count which was not further reduced in combination with azithromycin. Values represent means ± SEM. **p>0.01, ***p> 0.001 indicates CLP plus ceftriaxone or CLP plus (ceftriaxone + azithromycin) versus CLP.

## DISCUSSION

Macrolides are reported to produce immunomodulatory action by inhibiting production of reactive oxygen species, pro inflammatory cytokines and transcription factor of inflammation like nuclear factor kappa B (NF-κB) and activator protein 1 (AP-1). They are also known to inhibit adhesion of molecules on endothelial cells and infiltration of neutrophils from blood to tissues (11, 12). In a previous *in vitro* study, azithromycin produced dose dependent reduction in cytokines and nitrate/nitrite levels in LPS stimulated J774 macrophages (13). In another report, azithromycin had demonstrated protective effect from systemic damage induced by lethal dose of LPS in mice by inhibiting TNF-α (9). Beneficial effect of azithromycin on pulmonary inflammation has been established earlier by single oral or intraperitoneal dose (9, 14, 15, 16).

In the present study, we have demonstrated the survival benefit of azithromycin in murine model of sepsis. Clinically, sepsis is accompanied with infection and hence azithromycin was evaluated in CLP model of sepsis which correlates more with human sepsis. Azithromycin administered alone did not provide significant protection in CLP mice. This could be due to its inability to stop the progression of infection. For survival benefit in CLP model of sepsis, it is important to manage both infection and inflammation. Infection can be very well handled by treatment with suitable antibacterial agents. Therefore, in the current study we have combined azithromycin with ceftriaxone, a broad spectrum β-lactam antibiotic to combat infection. A sub-effective dose of ceftriaxone was determined earlier in CLP mice and further used in combination with immunomodulatory dose of azithromycin. Ceftriaxone plus azithromycin produced beneficial protective effect in CLP challenged mice compared to standalone group. This effect of combination was not associated with synergistic antibacterial activity of both as azithromycin did not potentiate the antibacterial effect of ceftriaxone as observed through the bacterial load in blood, lung and peritoneal fluid.

LPS and CLP challenge mice significant reduced the WBC count compare to normal mice possibly due to immunosuppression observed during later phase of sepsis. Similar decrease in leukocyte count was also noted in previous publications (17, 18). In both the models, azithromycin neither elevated nor further suppressed the WBC count, thus suggesting it to have no immunosuppressive property. Sepsis is reported to produce hypodynamic state by reducing body temperature and blood glucose (19). Therefore these parameters were measured in all the groups. As expected, mice subjected to LPS treatment or CLP challenge significantly reduced the blood glucose and body temperature while azithromycin treatment significantly reversed them.

In sepsis, reactive oxygen species and MPO production surpass antioxidant defenses and leads to a state of oxidative stress that triggers inflammation and causes direct mitochondrial damages, which results into sepsis-induced organ dysfunction (20). Increased free radicals that caused peroxidation of membrane lipids have been associated with reduction of GSH content (21). GSH an endogenous antioxidant, not only provides protection from reactive oxygen species but also inhibit the production of several inflammatory cytokines and chemokines (22). Unlike normal control group, LPS treated mice showed decreased in GSH while azithromycin significantly increased its levels. In our study, we have demonstrated elevation in plasma MPO in LPS treated mice which was significantly reduced by azithromycin. In the CLP mice, as expected there was decrease in plasma GSH and increase in MPO levels compared to sham group. Azithromycin in combination with ceftriaxone produced significant increase in plasma GSH and suppression of MPO levels.

Previous studies have demonstrated that TNF-α, IL-1β and IL-6 are the key contributors in the pathogenesis of sepsis (23, 24, 25). They are primarily responsible for apoptosis, necrosis of tissues and thereby multiple organ failure. TNF-α and IL-1β are reported to initiate inflammatory response in early sepsis and they stimulate the production of IL-6 which dominates in the later phase of sepsis (26). IL-6 has been extensively examined in patients with sepsis and its concentrations correlates more closely with severity and clinical outcome (26). Present study demonstrated elevated levels of these cytokines in the LPS and CLP challenged mice, while azithromycin significantly mitigated its levels. Sepsis associated lung injury is considered to be a leading cause of death (27). Azithromycin in combination with ceftriaxone produced statistically significant reduction in levels of inflammatory cytokines in lung tissue. Azithromycin treatment also led to a significant suppression of lung MPO activity, suggesting that azithromycin inhibited the neutrophil infiltration into the lung parenchyma and alveolar spaces. Finding of our study is in agreement with earlier report where azithromycin had demonstrated beneficial effect in combination with ceftriaxone in mouse model of lethal pneumococcal pneumonia due to its immunomodulatory activity (28). In the present study, we have demonstrated immunomodulatory activity of azithromycin in both LPS and CLP sepsis model. Azithromycin significantly reduced the inflammatory cytokines, MPO levels and increased the GSH levels which are otherwise altered to a pathological level in sepsis leading to organ failure. We have further demonstrated protective role of azithromycin in sepsis associated lung damage.

Findings of this study indicate that combination therapy of ceftriaxone and azithromycin has the potentials to improve survival in critically ill patients with bacterial infection.

## METHODS

All the experimental protocols were approved by the Institutional Animals Ethics Committee (IAEC) of Wockhardt Research centre, India. Female Swiss albino mice, weighing 25-30 gm were used for entire study. Mice were housed in cages with standard rodent feed and drinking water provided *ad libitum*. The light cycle was controlled automatically (on at 7:00 a.m. and off at 7:00 p.m.), and the room temperature was regulated to 18-22°C with 40-70% humidity.

### LPS induced sepsis model

The purpose of evaluating azithromycin in this model was to determine its immunomodulatory dose in our experimental set up. LPS derived from *Escherichia coli* serotype 0127:B8 (Sigma-Aldrich) was dissolved in saline and injected intraperitoneally (*i.p*.) in a fix volume of 0.25 mL to induce sepsis in mice. Two different LPS doses were used; a lethal dose of 1500 µg/mouse for survival study where monitoring was done up to 24 h and a sub-lethal dose of 1000 µg/mouse for estimation of body temperature, blood glucose, total white blood cell (WBC) count, cytokines, myeloperoxidase (MPO) and glutathione (GSH) levels.

### Treatment groups for LPS induced sepsis model

For survival experiment, 1 h prior to lethal dose of LPS, mice were administered vehicle (saline) and azithromycin (Azithral; Alembic Pharmaceutical) at doses of 25, 50 and 100 mg/kg, intraperitoneally. From this survival study, two effective doses of azithromycin were further selected to assess the different biochemical and physiological parameters in mice exposed to sub-lethal dose of LPS.

CLP induced sepsis model

Mice were anesthetized with mixture of ketamine/xylazine (100/10 mg/kg, *i.p*.) and abdomen was disinfected with iodine. A small mid-abdominal incision was made to expose the cecum. A distended portion of the cecum just distal to the ileocecal valve was isolated and ligated with a 3-0 silk suture in a manner not to disrupt bowel continuity. The ligated portion of cecum was punctured twice with an 18-gauge needle. To ensure the patency of the punctured portion, cecum was gently squeezed until small quantity of feces extruded through them. The cecum was placed back in the abdomen, and the incision was closed with sutures. The mice were then resuscitated with 1 mL of saline injected subcutaneously and returned to their cages with free access to feed and water. Sham-operated controls were treated in an identical manner, but without ligation and puncture. Survival was assessed twice a day for 5 days. In a separate study, parameters like body temperature, blood glucose, total WBC count, plasma GSH, cytokines and MPO levels in plasma and lung tissue along with bacterial count in blood, peritoneal fluid and lung homogenate were measured after 18 h of CLP challenge.

### Treatment group for CLP induced sepsis model

From the initial survival studies performed in CLP mice (data not shown), sub-effective dose of ceftriaxone (100 mg/kg, *s.c*.) providing protection in 37.5 % of mice was determined. The survival study included a sham group, CLP mice treated with vehicle (saline), ceftriaxone (100 mg/kg, *s.c*), azithromycin (100 mg/kg, *i.p*.) and both in combination. For estimation of biochemical and other parameters, groups included were sham control, CLP mice treated with vehicle (saline), ceftriaxone (100 mg/kg, *s.c*) and ceftriaxone (100 mg/kg, *s.c*) plus azithromycin (100 mg/kg, *i.p*.), all administered 3 h post CLP.

### Experimental protocol

For estimation of biochemical and other parameters, 16 mice were included in each group. Mice were made septic by treatment with either LPS or by CLP technique and 18 h later body temperature was measured. Half of the mice were bled through tail vein for glucose estimation and followed by blood collection through retro orbital sinus in EDTA tubes for total WBC count. Remaining mice were bled through retro orbital sinus in heparinised tubes to obtain plasma for estimation of cytokines, MPO and GSH. Lungs were harvested from all mice, rinsed with saline and weighed. Lungs from six mice were homogenized with chilled saline to obtain 20% homogenate from which 100 µl was used for bacterial count and the remaining was centrifuged at 15000 rpm for 10 minutes at 4 ºC for cytokine measurements. Lungs from remaining mice were homogenized with 50 mM potassium phosphate buffer (pH 6.0) containing 0.5 % hexadecyltrimethylammonium bromide, sonicated and centrifuged to obtain supernatant for MPO estimation.

### Measurements

#### Body temperature and blood glucose

Rectal temperature was measured 18 h after LPS and CLP challenge using digital thermometer (CareTouch^®^). Blood glucose was estimated using calibrated Bayer Contour^®^ TS glucometer.

#### Estimation of Total WBC count

Blood samples collected in EDTA were analyzed on automatic blood cell counter (Sysmex haematology analyzer) for determining the total white blood cells.

#### Estimation of Plasma cytokine levels

Tumor necrosis factor-α (TNF-α), Interleukin 6 (IL-6), and IL-1β were estimated using commercially available enzyme-linked immunosorbent assays (ELISAs), according to manufacturer’s instruction (R & D Systems Inc, USA).

#### Measurement of MPO activity in plasma and lung

MPO activity was determined by O-dianisidine method with modification for 96 well plate. The assay mixture consisted of 30 µL of 0.1M phosphate buffer (pH 6), 30 µL of 0.01M hydrogen peroxide (Merck), 20 µL plasma and lung supernatant, 170 µL deionised water and 50 µL of freshly prepared 0.02M of O-dianisidine (Sigma-Aldrich) solution. O-dianisidine solution was added last and absorbance was recorded per minute at 460 nm up to duration of 10 minutes using spectrophotometer (SpectraMax^®^ Plus 384 Microplate Reader). MPO activity was expressed in µM/min/mL of plasma and in U/gm of tissue.

#### Determination of glutathione (GSH) content

GSH was determined using Ellman’s reagent [5, 5’-Dithiobis (2-nitrobenzoic acid) or DTNB]. GSH (the most abundant non-protein thiol) reduces DTNB to form a stable yellow product (5-mercapto-2-nitrobenzoic acid), which can be measured colorimetrically. The reaction mixture consisted of 50 µL of clear plasma, 250 µL of 0.1M phosphate buffer (pH 6) and 25 µL of DTNB reagent (4 mg/mL dissolved in 1% sodium citrate). This mixture was then incubated for 10 minutes at 37 ºC and the absorbance was read at 412 nm using spectrophotometer. The GSH concentration was determined using a standard curve constructed with different concentrations of reduced L-glutathione (Sigma-Aldrich).

#### Evaluation of Bacterial Clearance

Bacterial load was determined in blood, lung homogenate and peritoneal lavage. In brief, mice were anesthetized with ketamine/ xylazine after 18 h of CLP. The blood samples were collected in heparinised tubes through retro orbital sinus, mixed and placed on ice bath. Mice were then injected 2 mL sterile saline intraperitoneally and abdomen was gently massaged. Later the skin over the abdomen was cleansed with 70% alcohol and cut open to expose the peritoneal cavity to collect the peritoneal lavage fluid. Peritoneal lavage fluids, blood and lung homogenate (100 µL) were placed on ice and serially diluted with sterile saline. 10 µL of each diluted samples were placed on trypticase soy agar plates (BD Biosciences, San Diego, CA) and incubated at 37°C for 24 h. The numbers of bacterial colonies were then counted and expressed as colony-forming units (CFU) per milliliter of blood and peritoneal lavage and CFU per milligram of lung tissue.

### Statistics

Survival data were analyzed using the log rank test and Fischer exact test. Data were represented as mean ± S.E.M. Differences among groups were assessed using one-way analysis of variance (ANOVA) test, followed by Dunnett’s multiple comparison post hoc test. Probability of p<0.05 was considered to be significant. All statistical analysis was performed using Graphpad prism statistical software (version 5).

## Acknowledgment

We thank Wockhardt for granting permission to perform the studies.

